# Uncovering locomotor learning dynamics in people with Parkinson’s disease

**DOI:** 10.1101/2024.10.17.618863

**Authors:** Aram Kim, Nicolas Schweighofer, Giselle M. Petzinger, James M. Finley

## Abstract

Locomotor learning is important for improving gait and balance impairments in people with Parkinson’s disease (PD). While PD disrupts neural networks involved in motor learning, there is a limited understanding of how PD influences the time course of locomotor learning and retention. Here, we used a virtual obstacle negotiation task to investigate whether the early stages of PD affect the acquisition and retention of locomotor skills. On Day 1, 15 participants with PD and 20 age-matched controls were instructed to achieve a specified level of foot clearance while repeatedly stepping over two different virtual obstacles on a treadmill. We assessed online performance improvement on Day 1 and overnight retention after at least 24 hours on Day 2. We used a hierarchical Bayesian state-space model to estimate the learning rate and the degree of interference between the two obstacles. There was a 93% probability that people with PD learned the locomotor skill faster than controls, but there was limited evidence of group differences in interference between the two heights of obstacles. Both groups improved their performance to a similar magnitude during skill acquisition and performed similarly during retention on Day 2. Notably, a slower learning rate was associated with greater online performance improvement, while lower interference was linked to better overnight retention, and this effect was strongest for the control group. These results highlight that people with early-stage PD retain the ability to use multisensory information to acquire and retain locomotor skills. In particular, our finding that people with early-stage PD learned faster than age-matched controls may reflect the emergence of compensatory motor learning strategies used to offset early motor impairments in people with PD.

## Introduction

Parkinson’s disease (PD) is a multifaceted, neurodegenerative disorder resulting primarily from the loss of dopaminergic neurons in the basal ganglia (1,2) and characterized by motor symptoms such as gait and postural impairments (3) and non-motor symptoms such as cognitive impairment (4–6). Dopamine replacement therapy is the gold-standard treatment to alleviate motor symptoms in PD (3) but often does not relieve instability during gait (3,7). A complementary approach to improve motor impairments not responsive to dopamine replacement therapy is motor skill learning (8–11), which promotes neuroplasticity and reduces motor impairments in people with PD (8–10).

Motor skill learning (12–15) engages different neural networks based on the features of the training task and across various phases of the learning process. Skill acquisition involves changes in the capacity to perform a skill and can be mediated by error-based learning, which engages cerebellar circuits (16–18), reinforcement of successful actions, which involves cortico-striatal circuits (19–21), or repetition (22–24) or explicit instructions (25–27), which both engage the prefrontal cortex. Retention refers to the maintenance of the capacity to perform acquired motor skills over time (28). When an individual learns multiple skills through interleaved practice, as is common in physical therapy, the performance of one skill may influence the performance of another. This influence may be in the form of interference, where the practice of a skill hinders performance in another (29,30), or facilitation, where the practice of one skill leads to improvements in the performance of another. Although interference may hinder initial skill acquisition, it can improve long-term retention (31). Understanding how Parkinson’s disease affects features of locomotor skill learning is critical for optimizing mobility-related rehabilitation.

Although PD leads to the degeneration of neural circuits involved in motor learning, there are mixed results regarding whether people with PD exhibit impairments in motor skill acquisition and retention. Some studies report that people with PD acquire motor skills more slowly than older adults (32–37), while most often, their learning rates are similar (38–48). Findings on motor retention also have been mixed; there is evidence that PD impairs retention (34, 38, 39, 44, 45) and that retention is comparable to controls (37,41,43,49,50). Further, there are varied results regarding the impact of interference during motor skill learning on retention in people with PD. Some studies suggest that interference during skill acquisition does not improve long-term retention in people with PD (35,47), while recent studies indicate that interference results in lower error and faster skill execution (51,52).

In addition, as most studies of motor learning in PD focus on tasks that rely on the upper extremities, we have only a limited understanding of how PD influences the learning of mobility-related skills. In one study that examined error-based learning during locomotion, people with PD learned to modify foot height during obstacle negotiation more slowly than age-matched controls (32,33). However, people with PD who performed a locomotor adaptation task adapted at similar (42,53) or even faster (54) rates than controls. Another limitation of most studies of locomotor learning in PD is that they often focus on single-day adaptive learning without examining how these skills are retained or how different skills interact during learning.

Here, we investigated how people with early-stage PD acquired and retained two potentially interfering virtual obstacle negotiation skills. Virtual reality provided a simple means to vary the task objectives by providing multiple obstacle heights and objectively quantifying performance. As initial skill acquisition primarily relies on error-based motor learning and cerebellar circuits, which are generally intact in people with the early stages of PD (39,55), we hypothesized that people with PD would acquire a locomotor skill at a similar rate to the controls. Similarly, we hypothesized that people with PD would have intact online performance improvement relative to the controls. However, switching between strategies or *set-shifting* requires frontostriatal circuits (56), which are impaired in people with PD (35,57). Therefore, we hypothesized that people with PD would have greater interference between obstacles than controls. Moreover, as PD impairs cortico-striatal circuits (57) and these circuits mediate the retention of motor skills (39,58,59), we hypothesized that people with PD would have impaired retention 24 hours after practice. Finally, we hypothesized that individual differences in locomotor skill learning would be associated with online performance improvement and overnight retention.

## Methods

### Participants

We recruited 37 individuals, including 15 people with PD from the neurology clinic at the University of Southern California and 22 age-matched adults as controls from the community (Table 1). Inclusion criteria for individuals with PD were: 1) no cognitive impairments, determined by two clinical experts using level-two criteria as recommended by the Movement Disorder Society task force (6,60), 2) ability to provide informed consent, 3) confirmed diagnosis of PD based on the UK Brain Bank criteria, 4) Hoehn and Yahr (H&Y) stage 1 to 3, and 5) stable on medication. Exclusion criteria for individuals with PD were 1) other neurological, cardiovascular, orthopedic, and psychiatric diagnoses that may affect walking, 2) Levodopa-induced hallucinations, and 3) freezing of gait. The inclusion criteria for the controls were 1) no cognitive impairments and 2) the ability to provide informed consent. Exclusion criteria for healthy older adults were neurological, cardiovascular, orthopedic, and psychiatric diagnoses. All participants with PD were tested on medication. We excluded one control for a technical issue and one control because they could not perform the virtual obstacle negotiation task. The Institutional Review Board at the University of Southern California approved study procedures, and all participants provided written, informed consent before testing began. All aspects of the study conformed to the principles described in the Declaration of Helsinki.

**Table 1.** Demographics and clinical assessments. Results are presented as the mean and standard deviation except for sex, Hoehn and Yahr (H&Y), and Disease duration. Sex and H&Y are presented as the number of participants. Disease duration is presented as the median and a range. Overground speed was calculated using the 10 Meter Walk Test (10MWT). Mini-BESTest: mini-Balance Evaluation Systems Test, MDS-UPDRS: Movement Disorder Society – Unified Parkinson’s Disease Rating Scales, LED: Levodopa equivalent dosage.

|  | **Controls (n=22)** | **PD (n=15)** |
| --- | --- | --- |
| **Age (years)** | 66 ± 10 | 65 ± 10 |
| **Sex** | 8 M, 14 F | 7 M, 8 F |
| **Treadmill speed (m/s)** | 0.93 ± 0.25 | 1.00 ± 0.24 |
| **Overground speed (m/s)** | 1.33 ± 0.29 | 1.35 ± 0.19 |
| **Mini-BESTest** | 23 ± 4 | 24 ± 4 |
| **MDS-UPDRS part III** | - | 20 ± 8 |
| **H&Y** | - | 1=2, 2=13 |
| **MDS-UPDRS total** | - | 39 ± 10 |
| **Disease duration (years)** | - | 3 (0 - 35) |
| **LED (mg)** | - | 520 ± 377 |

### Experimental Setup

Participants viewed a virtual environment through the head-mounted display from a first-person viewpoint while walking on a treadmill (Figure 1A). The virtual environment was developed using Sketchup (Trimble Navigation Limited, USA), and the participant’s interaction with the environment was controlled and recorded using Vizard (WorldViz, USA). The environment was displayed within the HTC Vive head-mounted display, which has a 110-degree field of view, a resolution of 1080 × 1200 pixels per eye, and a mass of approximately 550 g. The virtual environment consisted of a corridor with obstacles on the left and the right side (Figure 1B). Thirty-two virtual obstacles were placed along the corridor at random intervals between 4 m and 8 m. We used two different heights of obstacles to ensure that the skill was sufficiently challenging. In our pilot study, we found that a single height was learned too quickly, while most participants could not fully learn the task if given three obstacles. The final obstacle heights were 0.05 m (LOW) and 0.18 m (HIGH) (Figure 1E). Each obstacle height was associated with a different range of foot clearance that was defined as successful (0.05-0.09 m and 0.01-0.05 m for the LOW and HIGH obstacles, respectively). The depth of the obstacles was set to 0.10 m. Participants stepped over obstacles with the leg on the same side as the obstacle and movement in the mediolateral direction was prevented in the virtual environment. Participants wore a safety harness and lightly held onto handrails while walking (Figure 1A)

**Figure 1.**
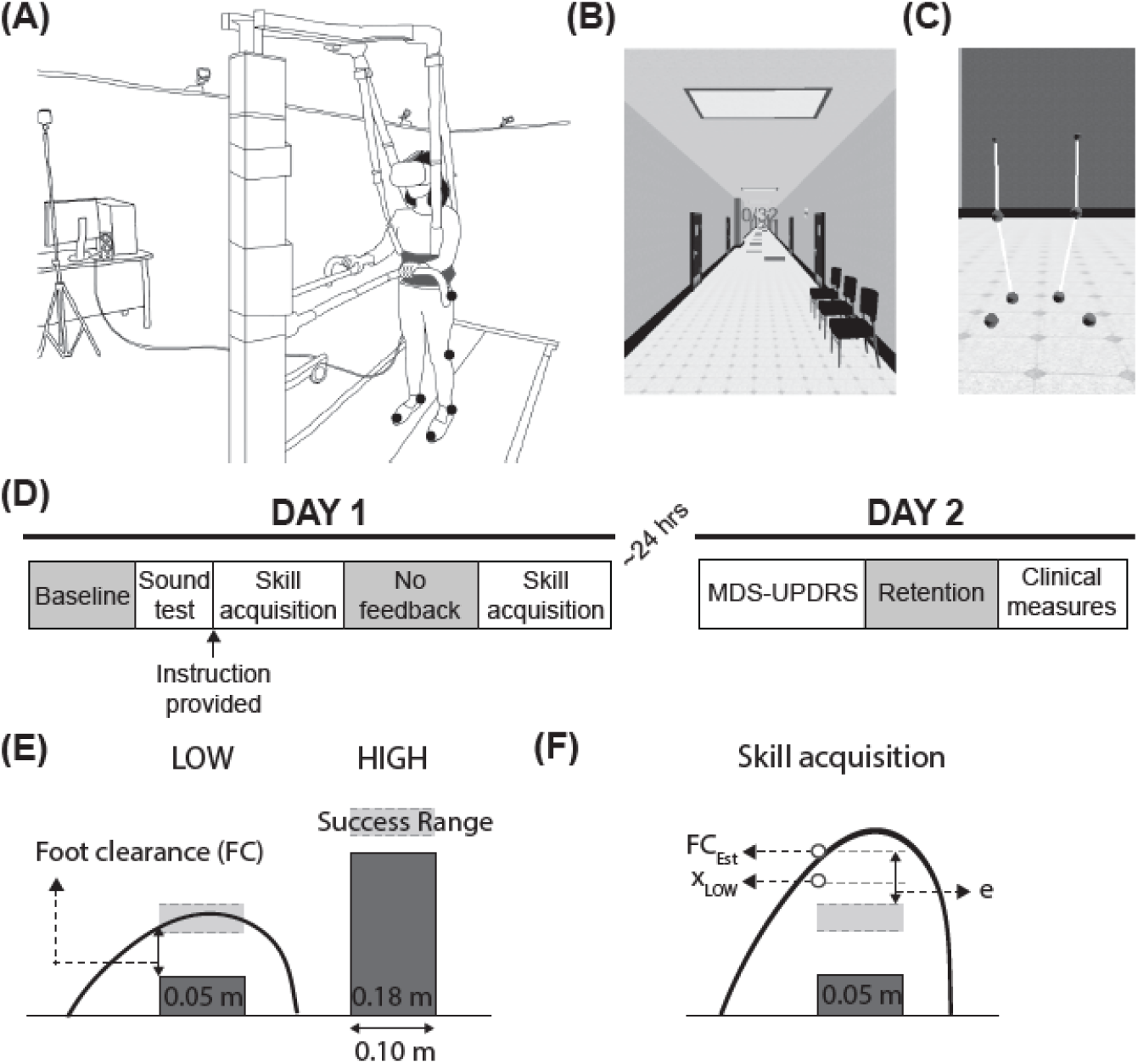
Experimental setup and protocol. (**A**) Schematic of the experimental setup. (**B**) The virtual environment. (**C**) Model of the lower extremities. Participants viewed the model from a first-person perspective. (**D**) Experimental protocol. MDS-UPDRS: Movement Disorder Society – Unified Parkinson’s Disease Rating Scale. (**E**) An example of foot trajectories and a definition of foot clearance (FC) for the LOW obstacle. Also, prescribed success ranges are shown in the shaded dotted box for both LOW and HIGH obstacles. (**F**) Schematic of state-space model variables for a LOW obstacle. Foot trajectories during skill acquisition are shown. FC_Est_ represents the motor output (estimated foot clearance) of the state-space model; ***x***_***L****OW*_ represents the estimated threshold of the success range for the LOW obstacle. In this example, as ***x***_***L***OW_ is higher than the success range, ***x***_***L***OW_ represents the estimated upper threshold; and e represents a motor error between the success range threshold and FC_Est_.

The velocity of the scene and the treadmill speed were synchronized using Python-based code in Vizard, and an inertial measurement unit within the head-mounted display controlled the orientation of the viewpoint. Participants’ lower extremities were presented in the first-person viewpoint, consisting of spheres at the toe, heel, knee, and hip on each side and line segments that connected the spheres (Figure 1C).

### Experimental Protocol

On Day 1, participants completed a novel, virtual obstacle negotiation task adapted from a prior study (33) on the treadmill (Bertec Fully Instrumented Treadmill, USA) at their self-selected speed (Figure 1D). During the baseline trials, we instructed participants to step over the obstacles as naturally as possible and to avoid collisions. If a collision occurred, participants heard an unpleasant sound. The baseline was divided into two trials for LOW and HIGH obstacles, respectively.

After completing the baseline, we instructed participants: “*You will step over obstacles of two different heights. Each obstacle has a different success range. This success range is invisible to you, so you will explore to find this range and maintain your foot within the success range while crossing. You will receive auditory feedback according to your performance*” (Figure 1D). Then, participants listened to examples of the three types of auditory feedback: 1) a pleasant sound if the foot clearance was within the success range, 2) an increment sound if the foot clearance was lower than the success range, indicating the need to increase foot clearance on the next obstacle with the same height, and 3) a decrement sound if the foot clearance was higher than the success range, instructing them to decrease foot clearance on their next obstacle with the same height. The duration of the sound was scaled to the distance from the foot and the success range during the crossing. After participants listened to all possible sounds, they were asked to identify 30 sound examples (*sound test*), including the pleasant sound, the unpleasant sound, two increments, and two decrement sounds that differed in duration (100% and 40% duration). If participants achieved less than 80% accuracy, they took another test with the same set of possible sounds until they achieved at least 80% accuracy. After they passed the sound test, participants started the skill acquisition trials.

Participants practiced stepping over a total of 192 obstacles in six 32-obstacle bouts. Auditory feedback was provided according to the performance during skill acquisition trials, and a score representing the number of successful obstacles was presented on the top of the virtual environment (Figure 1B). We tested *online performance improvement* during the second to last bout by removing all auditory feedback other than collision feedback (Figure 1D). The instruction was the same as in other practice bouts. For the last bout, participants received performance feedback again (Figure 1D).

After approximately 24 hours, participants revisited the laboratory to complete *overnight retention* trials (Day 2) (Figure 1D). During the retention test, participants received no auditory feedback regarding their performance other than collisions. We also assessed the symptom severity of PD using the Movement Disorder Society – Unified Parkinson’s Disease Rating Scales (MDS-UPDRS) and gait and balance using clinical assessments. These assessments included a 10-meter walk test (10MWT) and Mini-Balance Evaluation Systems Test (Mini-BESTest).

### Data Acquisition

A 10-camera Qualisys Oqus motion capture system (Qualisys AB, Sweden) captured position data at 100 Hz from reflective markers placed on the following landmarks on participants’ lower extremities: second toes, heels, lateral epicondyle of the femur, and greater trochanter.

### Data analysis

Foot clearance was defined as the minimum foot distance from the top of the obstacle (Figure 1E) and was calculated using the foot trajectory in the sagittal plane when the foot was passing over the obstacle in real time during experiments. Performance error was calculated as the difference between foot clearance and the closest success range threshold. For instance, if foot clearance was lower than the success range, performance error was the difference between foot clearance and the lower success range threshold. We computed online performance improvement as the difference in performance error between no feedback trials and baseline, with more negative values corresponding to greater reductions in error. We then defined overnight retention as the difference in performance error between the retention trials on Day 2 and the end of practice on Day 1. These values were generally positive, with larger values corresponding to greater forgetting between Days 1 and 2.

#### State-Space Model of Skill Acquisition

We used a single-state model to capture how the foot clearance (FC) for each obstacle(n) for each participant (p). This model contains a single state representing the threshold of the success range for each obstacle height 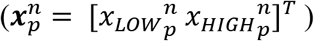 and the motor output, which is estimated foot clearance (*FC*_*Est*_)(61) (Figure 1F). These variables are updated as follows.

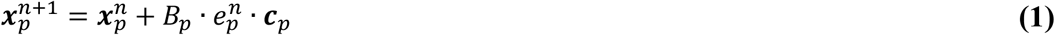

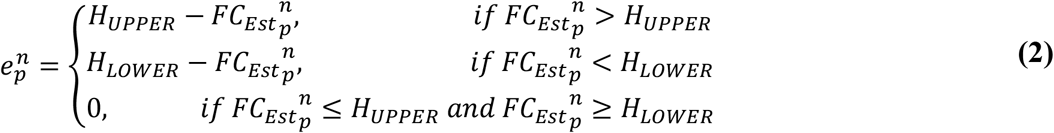

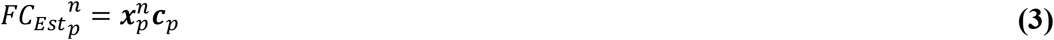

0 ≤ B_*p*_ ≤ 1 is a learning rate, which is the rate of the reduction in subsequent foot clearance for an error 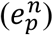 such that a B_*p*_ value of 1 would correspond to a reduction in foot clearance equal to the size of the error. The context variable is ***c***_*p*_ = (1 q_*p*_)^*T*^ for LOW or ***c*** = (q_*p*_ 1)^*T*^ for HIGH obstacles that captured how much an error on an obstacle of a given height influenced the state estimate of the other height obstacle. The parameter 0 ≤ q_*p*_ ≤ 1 modulated the level of interference such that a q_*p*_value equal to 1 corresponded to full interference (Eqn. 1). 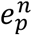 was calculated by the difference between 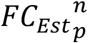 and success range of the obstacles (*H*) (Eqn. 2). When 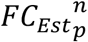 was within the threshold (*H*), the 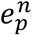 is zero; otherwise, it is given by the difference between the current foot clearance and the height of the given upper (*H*_*UPPER*_) or lower (*H*_*LOWER*_) threshold (success range). The state vector 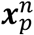 was passed and scaled by ***c***_*p*_ to estimate 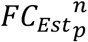 (Eqn. 3).

#### Hierarchical Bayesian Estimation of Model Parameters

Hierarchical Bayesian estimation relies on Markov Chain Monte Carlo analysis to estimate the posterior distributions of the individual parameters for each participant and the sample population as a whole (62). This form of estimation reduces the overall variance of the model by using the full dataset to estimate model parameters instead of fitting each participant’s data separately. This analysis also shifts the focus of our inference to parameter uncertainty instead of point estimates of the values of the model parameters (63). This model had three levels: an individual level (level 1), a participant level (level 2), and a population level (level 3).

##### Level 1: Modeling intra-individual variations

The observed food clearance 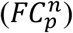 was modeled with a generalized Student’s t-distribution to account for outliers.

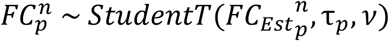

where 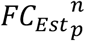 is the center of the distribution and τ_*p*_ is a participant-specific scale parameter, and *τ* is the degree of freedom.

##### Level 2: Modeling inter-individual variations

To account for the differences between individuals within each group (g), we modeled random variables following different probability distributions for each participant for the parameters. Thus, the individual parameters *B*_*p*_, q_*p*_, and τ_*p*_ were modeled with the following prior distributions.

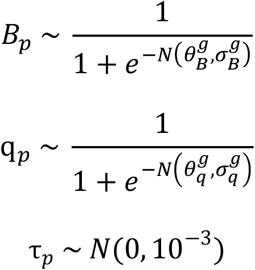

*N*(*µ, σ*) is the normal distribution with a mean *µ* and standard deviation *σ*. Learning rate *B*_*p*_ and interference parameter q_*p*_ are constrained between 0 and 1 via the sigmoid function. τ_*p*_ was sampled from an uninformative normal distribution. We also initialized 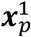 as free parameters that were sampled from participant-specific informative prior distributions from *FC*_*B*a*s*eline_ for LOW and HIGH obstacles separately and scaled by a scaling factor (*k*_*p*_), which were non-negative with truncated normal distributions.

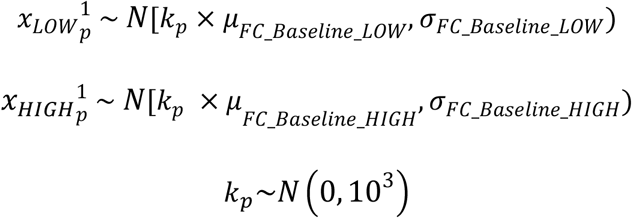

##### Level 3: Modeling the population level

The hyper-parameters 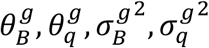, *and τ* govern the prior distributions of the individual parameters in level 1 or 2. We used normal priors for the location hyper-parameters *θ* and inverse-gamma priors for the scale hyper-parameters *σ* and *τ*. All hyper-parameters were initially sampled from uninformative prior distributions so that the posterior distributions were primarily influenced by the data (Table 2).

**Table 2.** Participant-dependent parameters and population-based hyper-priors in the best model.

| Model parameters | Priors | Hyper-priors |
| --- | --- | --- |
| Learning rate | $B_p \sim \frac{1}{1 + e^{-N(\theta_B^g, \sigma_B^g)}}$ | $\theta_B^g \sim N(0, 10^3)$<br>$\sigma_B^{g^2} \sim 1/\Gamma(10^{-3}, 10^{-3})$ |
| Interference | $q_p \sim \frac{1}{1 + e^{-N(\theta_q^g, \sigma_q^g)}}$ | $\theta_q^g \sim N(0, 10^3)$<br>$\sigma_q^{g^2} \sim 1/\Gamma(10^{-3}, 10^{-3})$ |
| Degrees of freedom | $\nu$ | $\nu \sim 1/\Gamma(10^{-3}, 10^{-3})$ |

### Bayesian Parameter Estimation

We used hierarchical Bayesian estimation with Markov Chain Monte Carlo (MCMC) to estimate individual- and population-level parameters (62) from the state-space model for all participants. MCMC incorporates the prior distributions and measured data to generate a posterior probability distribution of each parameter based on Bayes’ rule (63). We estimated parameters using three parallel chains with 20,000 iterations in each chain, 10,000 iteration burn-in, and thinning of 10 for three chains, totaling 3000 samples in *Rjags* (v4-10) in R (version 3.5.1). We verified the MCMC convergence using 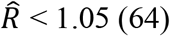 (64) for all parameters in the model. The low 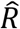 indicates that the three parallel chains are well-mixed, and all chains are converged to the target posterior distribution. In addition, we checked rank-normalized 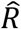 as a visual criterion (65) to ensure that MCMC searched a similar sampling space across three chains and the final results were similar regardless of the initial sample. If the search was done in a similar sampling space, rank normalized 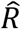 displays a uniform distribution. For all parameters, we visually verified the uniformity. Finally, we used functions from rjags to generate summary statistics and posterior distributions for the parameters in our model. Model comparisons were performed using the Watanabe-Akaike Information Criterion (WAIC) comparing the full model with both parameters (*B*_*p*_ and q_*p*_) and simpler models with each parameter. We confirmed that the full model with parameters *B*_*p*_ and q_*p*_ was the most favorable to explain our observed data.

### Bayesian Hypothesis Testing

To determine if there was a credible difference in learning rate or interference between groups, we transformed the posterior probability distributions of the population parameters using a sigmoid function. We then computed the difference between these distributions and discarded the proportion falling outside the 95% credible interval. Next, we calculated the proportion of the remaining distribution above and below zero with the probability of direction ranging from 50% to 100%. The greater of these two proportions represented the probability of direction. A bias in the posterior distribution toward negative values indicates that the learning parameter in the PD group is greater than in the control group and vice versa. Additionally, a probability of direction greater than 95% provides strong evidence of a directional effect (66,67).

We also tested whether the difference between groups in learning rate and interference was practically meaningful using the Region of Practical Equivalence (ROPE) testing (63,68). ROPE testing involves defining a range of “dead zones” where the effect is negligible or too small to be of any practical relevance. If 95% posterior probability falls completely outside of the ROPE (i.e., probability outside of the ROPE is 95%), there is strong evidence in favor of the effect of dependent variables. On the other hand, if 95% posterior probability falls completely inside of the ROPE, there is strong evidence in favor of the null hypothesis. We computed the effect size of population parameters of learning rate and interference using their untransformed posterior distribution 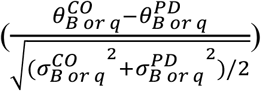. We used the ROPE of -0.1 and 0.1 as suggested by previous literature (68).

### Statistical Analysis

We first tested for the presence of demographic differences using two-sample independent t-tests. Moreover, we assessed differences in performance error during baseline between the groups using two-sample independent t-tests to confirm that initial performance was not different between the two groups. We also tested the effect of group and obstacle height on online performance improvement and overnight retention using repeated measures analysis of variance (rmANOVA). The dependent variable was performance error for each time point and each obstacle height. The independent variables were group (controls vs. PD), the height of obstacles (LOW vs. HIGH), and trials (baseline vs. no feedback for online performance improvement and end of practice vs retention on Day 2 for overnight retention). We assessed the main effects of and interactions between the independent variables. Finally, we used a linear mixed-effects model to determine whether individual differences in learning parameters (learning rate and interference) during skill acquisition predicted online performance improvement and overnight retention of the locomotor skill. The dependent variable was performance at each trial (no feedback or retention), and the independent variables were learning rate, interference, and group for each trial. We included interactions between each learning parameter and group to test a group-specific prediction. Further, as shown in our previous study (69), we assessed whether the individual difference in model-based final performance (error at the end of skill acquisition) predicted overnight retention. All statistical analyses were done in MATLAB R2020b (Natick, MA).

## Results

### Fitting the hierarchical Bayesian state-space models

Control participants and participants with PD reduced foot clearance error with practice (Figure 2), and this change in performance was captured well by the state-space model (Figure 3). Performance typically plateaued within 50-100 obstacles for all participants. The state-space model achieved good convergence with R-hat < 1.05 for all parameters and had an average RMSE of 0.03 ± 0.01 m, within the bounds of the targeted success range of 0.04 m.

**Figure 2.**
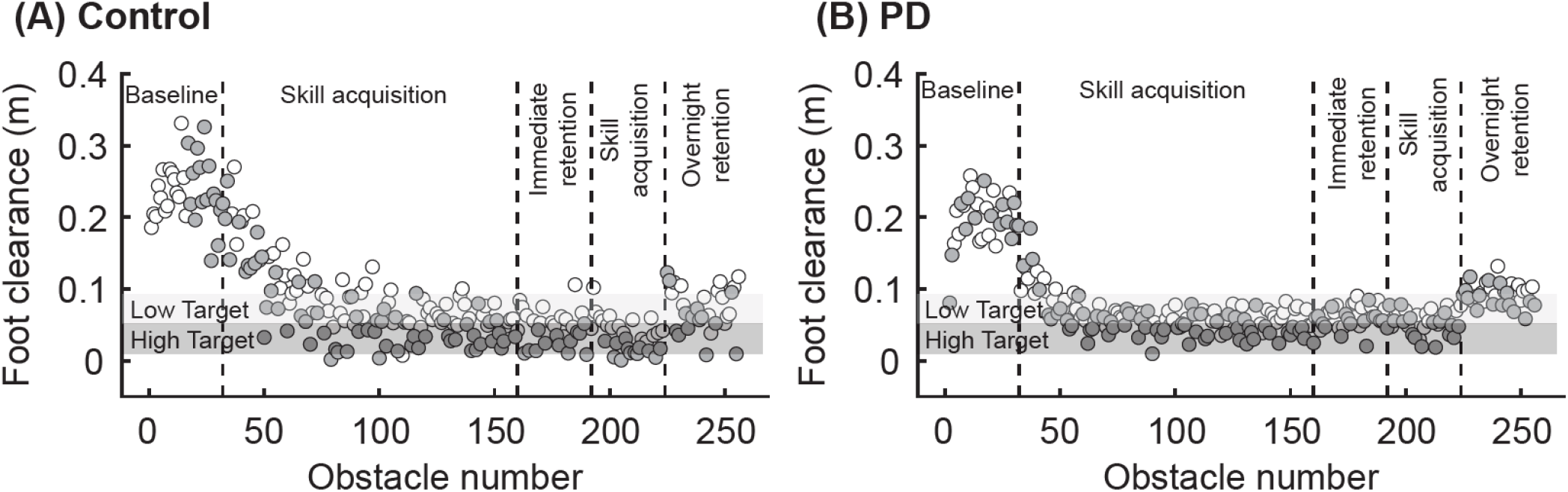
Example foot clearance data for (**A**) a control participant and (**B**) a participant with PD. Each data point represents foot clearance on a different obstacle. Empty circles correspond to LOW obstacles, and gray circles correspond to HIGH obstacles. The light and dark gray shaded areas represent the target range for the LOW and HIGH obstacles, respectively.

**Figure 3.**
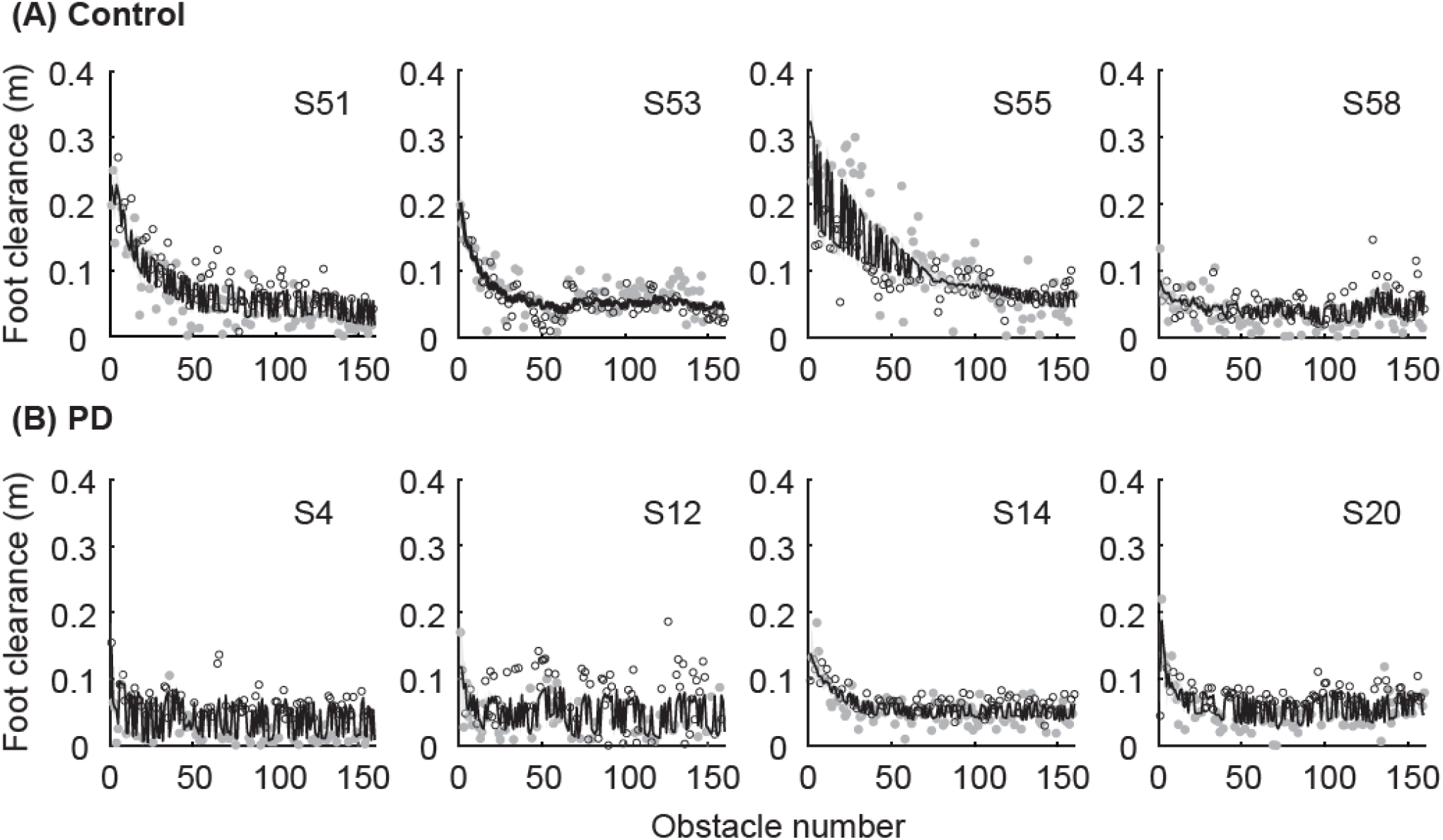
Observed foot clearance data during skill acquisition and model fit. Example participants (A) in the control group and (B) in the PD group. Data points represent the observed foot clearance as a function of obstacle number (Empty circle: LOW and Gray circle: HIGH). The solid black lines represent the model fit. The shaded areas represent 95% credible intervals, but these areas are frequently difficult to visualize as they are quite narrow.

### Motor Skill Acquisition

Using Hierarchical Bayesian estimation, we found a 93% probability that people with PD learn the skill faster than the controls (Figure 4). The median learning rate for the controls was 0.08 [95% CI: 0.05 – 0.14], and for people with PD was 0.14 [95% CI: 0.08 – 0.23] (Figure 4A). Estimates of the population difference in learning rate (the probability of direction) showed a bias toward a faster learning rate in people with PD than the controls (B_Group_ median and 95% CI: -0.06 [-0.15 0.02]), with 93% of the posterior probability distribution falling below zero. This difference between groups was also meaningful, as the effect size of the difference had a 93.1% probability of being outside of the ROPE [-0.1 0.1].

**Figure 4.**
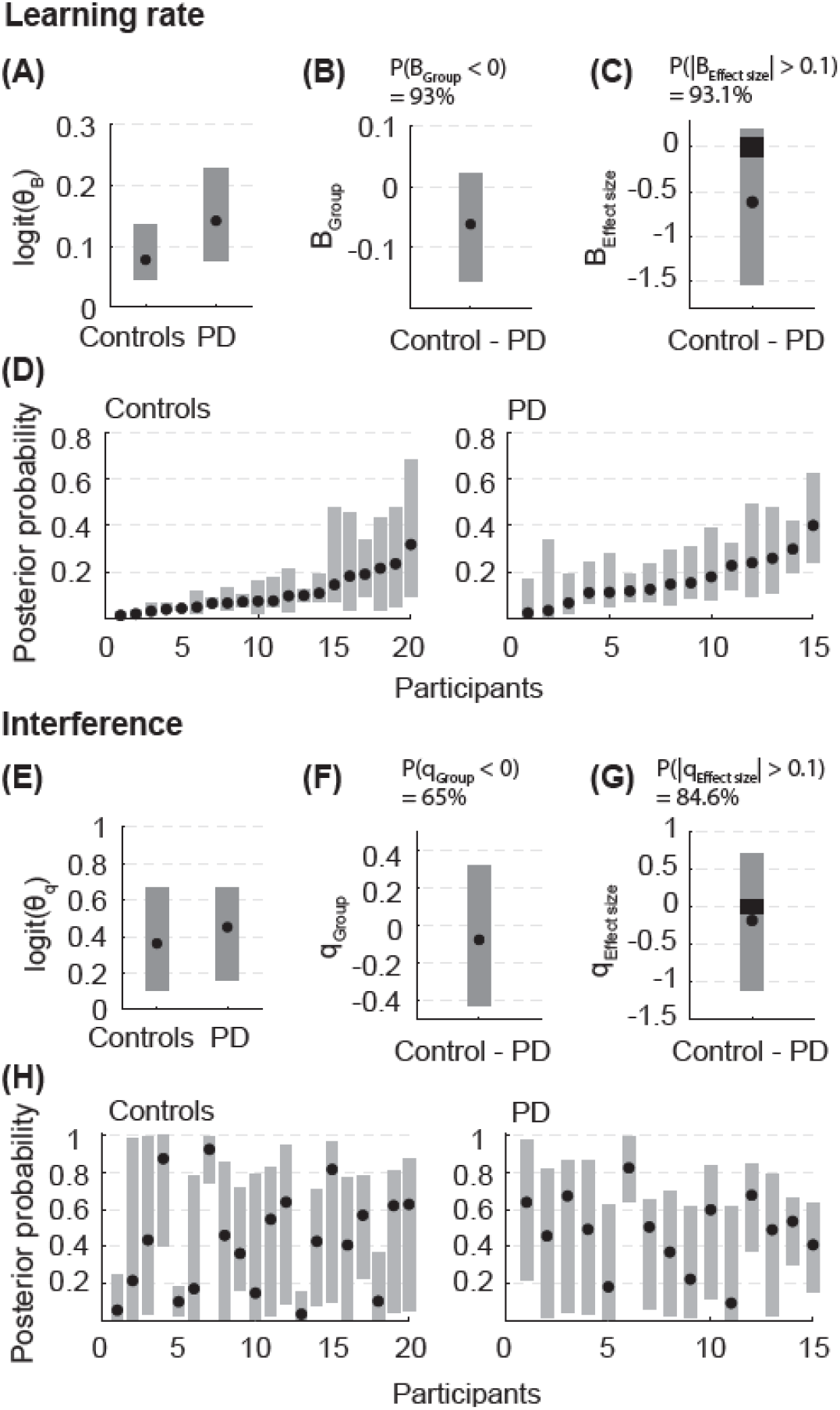
Posterior probabilities from the Bayesian estimation. Population 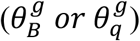 and individual parameter posterior probabilities for (**A-D**) learning rate and (**E-H**) interference. (**A** and **E**) The population parameter posterior probabilities are transformed using a sigmoid function for visualization. (**B** and **F**) B_Group_ and q_Group_ represent the difference in population parameters between the control and PD groups. The difference was calculated after transforming the posterior probabilities using a sigmoid function. P(group difference < 0) indicates the proportion of the group difference probability below zero. (**C** and **G**) B_Effect size_ and q_Effect size_ represent the effect size of the difference in population parameters between the control and PD groups. The effect size was calculated using untransformed posterior probabilities. P(|effect size| > 0.1) indicates the proportion of the probability outside the ROPE using the posterior probability effect sizes. The black rectangle represents the “dead zone” where the effect is negligible or too small to be of any practical relevance. (**D** and **H**) Individual learning rate and interference posterior probabilities were sorted according to the median learning rate for visualization purposes. For all panels, the black dots represent the median of the posterior probability, and the bars represent the 95% credible intervals.

However, we found limited evidence of the group difference in interference. There was only a 65% probability that interference between the LOW and HIGH obstacles was greater in people with PD compared to the controls (Figure 4). Given that the probability of direction ranged from 50% to 100%, this 65% probability of difference suggests limited evidence of the group difference. The median interference value for people with PD was 0.45 [95% CI: 0.16 – 0.66], and for controls was 0.36 [95% CI: 0.10 – 0.66] (Figure 4B). The posterior probability of difference spanned nearly equally in both directions (q_Group_ median and 95% CI: -0.08 [-0.43 – 0.31]). The uncertainty was likely due to large inter-individual variability in interference for both groups of participants. While several participants exhibited minimal interference (e.g., Control 1, 5, 13), some had interference values close to 1, indicating that errors experienced on obstacles of one height influenced subsequent foot clearance on obstacles of either height (e.g., Control 7, PD 6). Other participants showed large uncertainty in their interference with their CI, which spanned nearly the entire range. Lastly, the group difference in interference showed no practically meaningful effect, as only 84.6% of posterior probability fell outside of the ROPE.

### Online performance improvement

The practice of the virtual obstacle negotiation task led to reductions in errors that were maintained in the absence of auditory feedback. We used a rmANOVA to test for the main effects of the trial (Baseline on Day 1 vs. No feedback on Day 1), obstacle height (LOW vs. HIGH), and group (Controls vs. PD) as well as their interaction on performance error. We found a main effect of the trial (F(1,32) = 58.42, p < 0.001) on performance error, indicating that both groups had less error during no feedback relative to baseline on Day 1 (Controls: Δ = -0.08±0.08 m, PD: Δ = -0.07±0.05 m). We also found a main effect of obstacle height (F(1,32) = 6.94, p = 0.013) such that there was greater error for the HIGH obstacles than LOW obstacles (LOW: 0.06±0.06 m, HIGH: 0.09±0.06 m) (Figure 5). However, there was no main effect of group (F(1,32) = 1.75, p = 0.196) nor interactions between group and trial (F(1,32) = 0.01, p = 0.913), obstacle height and trial (F(1,32) = 1.01, p = 0.323), or group and obstacle height (F(1,32) = 1.02, p = 0.321) on performance error.

**Figure 5.**
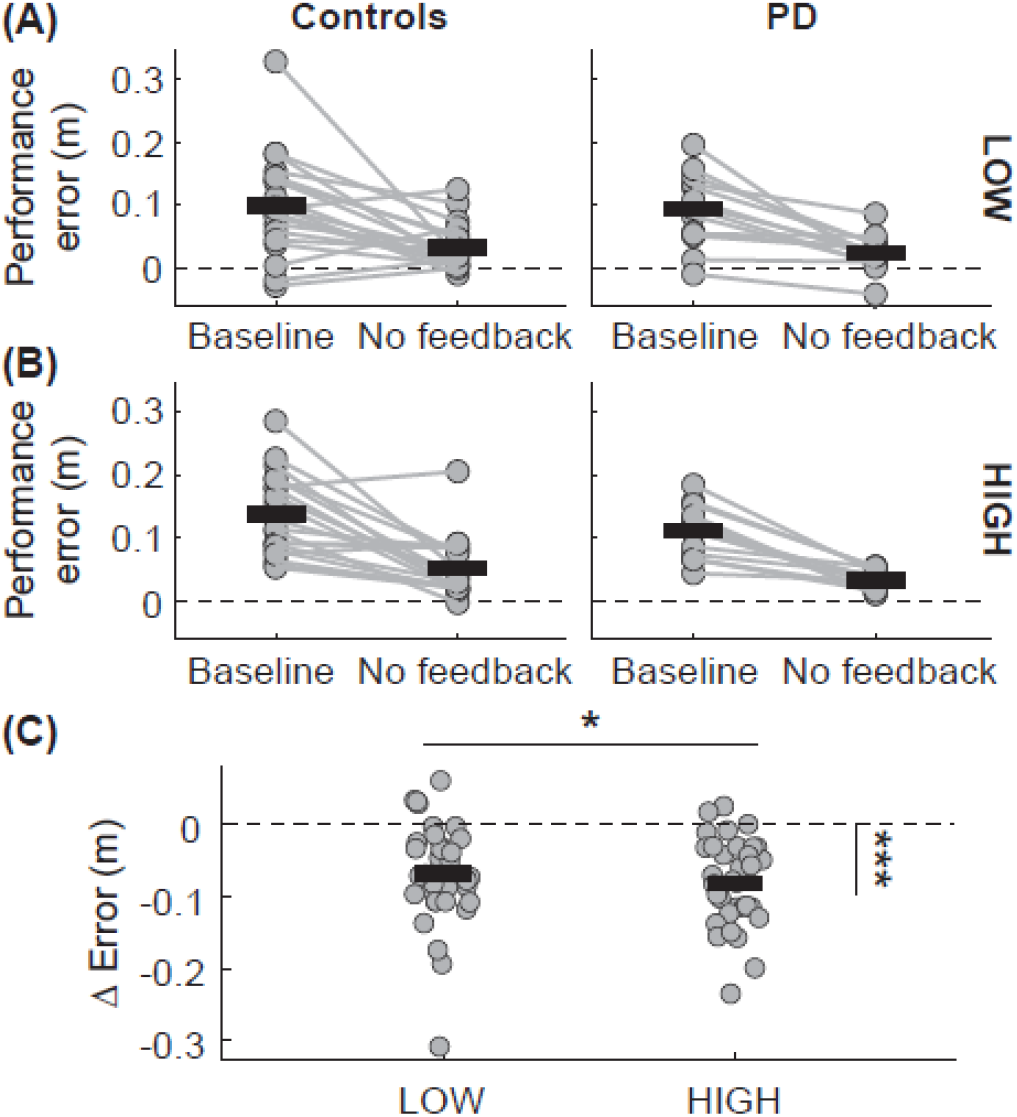
Performance error during baseline and no feedback trials. Performance error for each group (**A**) for LOW obstacles, (**B**) HIGH obstacles, and (**C**) Δ performance error (no feedback – baseline) for LOW and HIGH obstacles. The dots represent participants, and the connecting lines represent changes in performance error. Solid black lines represent means for each group. The dotted lines represent zero. Asterisks indicate statistical significance: *: p < 0.05, ***: p < 0.001.

### Overnight Retention

When participants returned 24 hours later, there was a significant increase in their performance error relative to the end of practice (End of practice vs. Retention, F(1,33) = 35.16, p < 0.001, Figure 6), indicating that both groups exhibited a loss of the skill from the end of skill acquisition (Controls: Δ = 0.04±0.04 m, PD: Δ = 0.03±0.02 m). However, the retention error was still significantly lower than their baseline error (rmANOVA F(1,33) = 47.89, p < 0.001). Moreover, there was a main effect of obstacle height (F(1,33) = 8.58, p = 0.006, Figure 6), such that there was a greater error for the HIGH obstacles than LOW obstacles (LOW: 0.03±0.03 m, HIGH: 0.05±0.04 m). However, there was no main effect of group (F(1,33) = 0.11, p = 0.743) nor interactions between group and trials (F(1,33) = 0.18, p = 0.678), obstacle height and trials (F(1,33) = 1.33, p = 0.257), or group and obstacle height (F(1,33) = 1.26, p = 0.27) on performance error. Overall, this suggests that both groups partially lost the locomotor skill after 24 hours.

**Figure 6.**
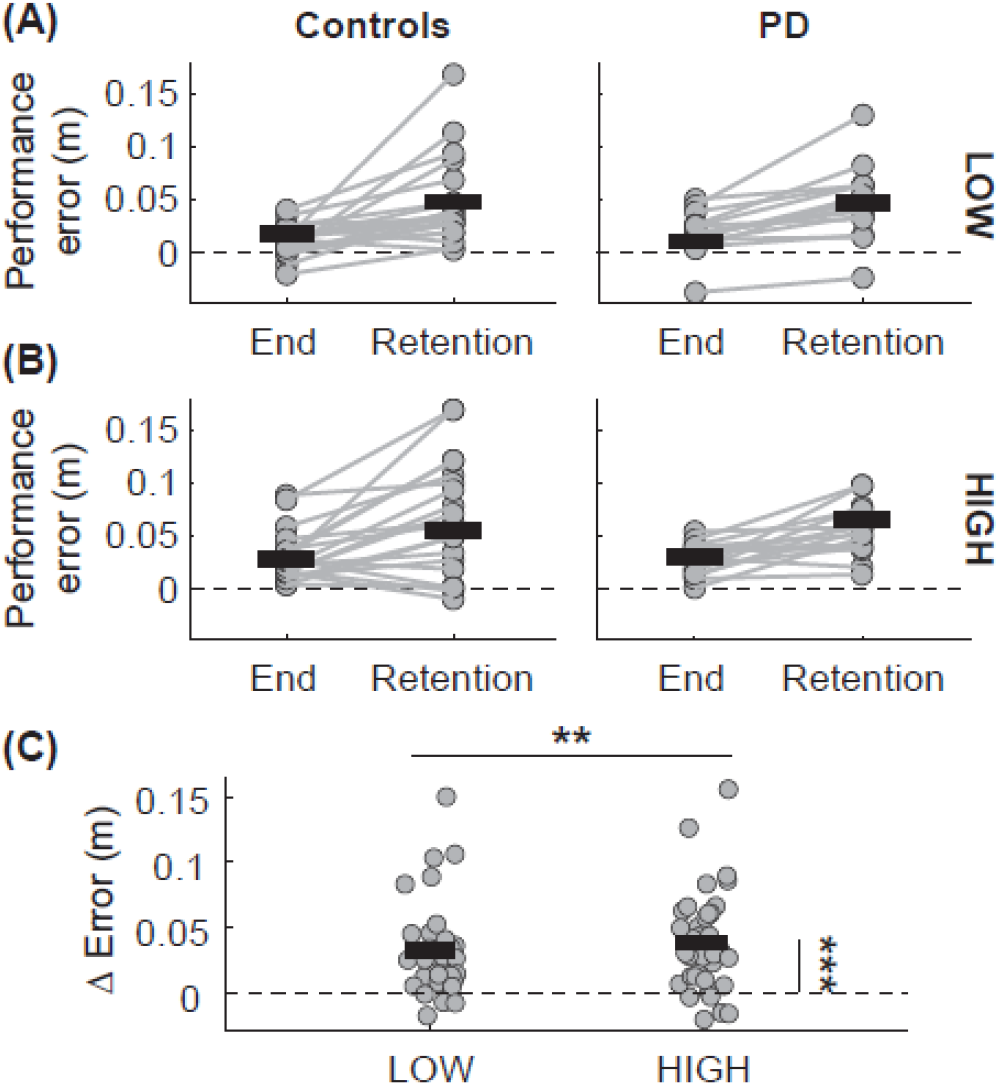
Performance error during the end of practice on Day 1 and retention on Day 2. Performance error for each group (**A**) for LOW obstacles, (**B**) HIGH obstacles, and (**C**) Δ performance error (retention – end of practice) for LOW and HIGH obstacles. The dots represent participants, and connecting lines represent changes in performance error. Solid black lines represent medians for each group. The dotted lines represent zero. Asterisks indicate statistical significance: **: p < 0.01, ***: p < 0.001.

### Associations between learning-related model parameters and learning performance

Online performance improvement and overnight retention were each associated with different learning-related parameters (Figure 7 A, B). There was a significant interaction between the group and learning rate on online performance improvement (t(28) = 2.27, p = 0.031, adjusted R^2^ = 0.58). Specifically, there was a more positive association between learning rate and online performance improvement in the controls than in the PD group (Figure 7A). Moreover, for overnight retention, the association between overnight retention and interference was different between groups (t(29) = 2.12, p = 0.043, adjusted R^2^ = 0.62). Specifically, the association was more positive in the controls than in the PD group, such that greater interference indicated worse overnight retention in the controls (Figure 7A).

**Figure 7.**
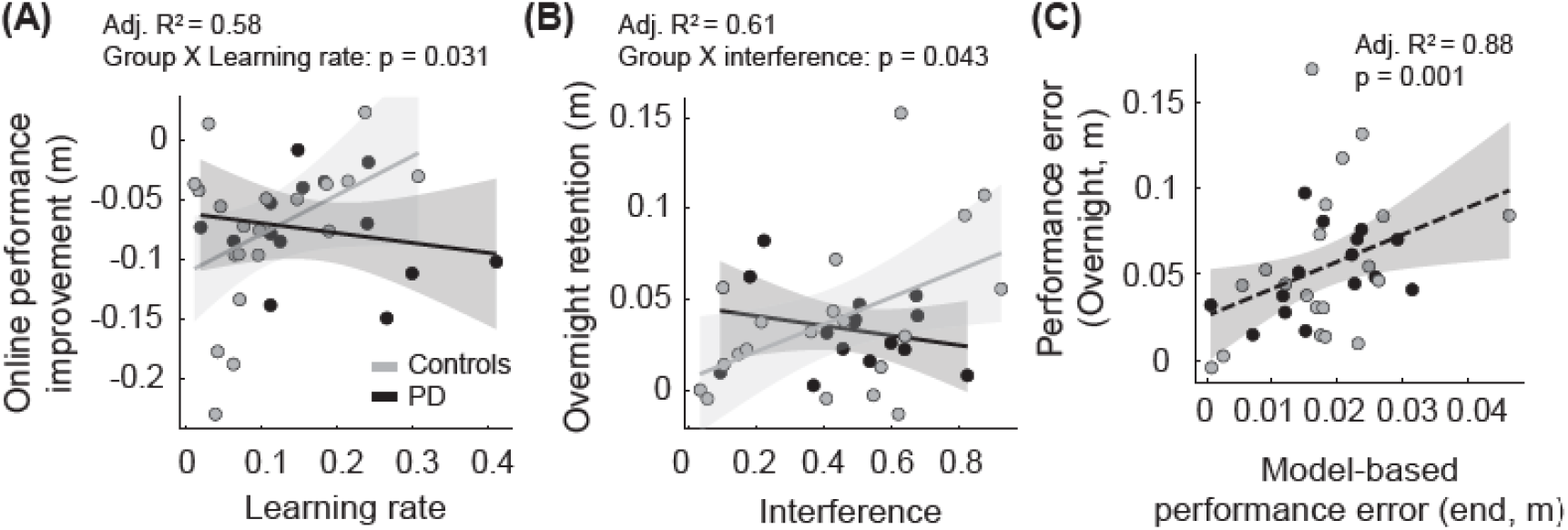
Associations between skill acquisition on Day 1 and learning performance. (**A**) Relationship between Δ performance error for online performance improvement (no feedback – baseline) on Day 1 and learning rate estimated by the Hierarchical Bayesian estimation. (**B**) Relationship between Δ performance error for overnight retention (retention – end of practice) on Day 2 and interference estimated by the Hierarchical Bayesian estimation. (**C**) Relationship between performance error during retention on Day 2 and model-based performance error at the end of skill acquisition estimated by the Hierarchical Bayesian estimation. For panels A and B, the black dots and lines represent the PD group data and regression fits, while the gray dots and lines represent the control data and regression fits. The dotted black line in (C) represents a regression fit for the full dataset.

In addition, overnight retention on Day 2 was associated with locomotor performance at the end of skill acquisition on Day 1 (Figure 7C). Due to the high variability in foot clearance during skill acquisition, we used performance error derived from the Hierarchical Bayesian estimation at the end of acquisition. We found that there was a significant association between overnight retention and model-based performance error at the end of skill acquisition (t(62) = 3.38, p = 0.001, adjusted R^2^ = 0.88), indicating that the final performance on Day 1 was indicative of overnight retention on Day 2.

## Discussion

Locomotor skill learning mediates improvements in mobility in response to gait interventions. Therefore, understanding the factors that influence locomotor skill learning is essential to inform the design and development of gait interventions for people with PD. Using a virtual obstacle negotiation learning task, we found that there is a 93% probability that people with PD acquire locomotor skills more rapidly than the controls. However, we found limited evidence of a group difference in interference as there is only a 65% probability that people with PD show greater interference between obstacles than the controls. People with PD also improved their locomotor performance and retained locomotor skills overnight relative to the controls. Lastly, learning rate and interference were associated with online performance improvement and overnight retention, respectively, but more strongly in the controls, and the final performance error on Day 1 was indicative of overnight retention regardless of the groups.

The evidence that people with PD acquire locomotor skills faster than age-matched controls may reflect what others report during the early stages of PD as compensatory hyperactivity in the cerebellum (55,70–74). The cerebellum processes sensorimotor error signals during motor learning (75–77) and has been shown to increase activity in response to dopamine depletion and disrupted cortico-striatal circuits. This hyperactivity is thought to help maintain better motor functions (71) and facilitate various types of learning, such as rule learning (78) and associative learning (79). Moreover, the cerebellum coordinates locomotor and balance control (80) and updates predictive motor commands in response to error signals (77). Given the cerebellum’s role, we speculate that this hyperactivity in the early stages of PD might enable people with PD to acquire error-based locomotor skills with auditory feedback more rapidly than age-matched controls. Future neuroimaging research is required to confirm our speculation. However, note that the proportion of the probability of direction did not reach 95%. This indicates that there is still inconclusive evidence for group differences in learning rate.

Moreover, the 93.1% probability outside of the ROPE suggests that there may be evidence for practical relevance, although the probability outside of the ROPE did not reach a practical relevance of 95%. The overall inconclusive evidence may be due to the small sample size to fully account for inter-individual variability. Future studies with larger samples should confirm how the learning rate is altered in people with early stages of PD compared to age-matched controls.

Contrary to our hypothesis, we also found that at the group level, both groups may face similar challenges in acquiring locomotor skills for obstacles of different heights, with only a slight tendency towards greater interference in the PD group than the control group. The probability of direction in interference was well below the 95% threshold typically considered necessary to establish a credible difference. Additionally, the probability outside the ROPE was 84.6%, indicating that the group difference may not be practically significant. There are two potential explanations. First, the large variability in posterior probabilities of interference may indicate that participants used different strategies to acquire the skills for various obstacles. For instance, the credible intervals for most participants spanned nearly the entire range of the posterior probability, indicating that the degree of interference could vary widely. This suggests that individuals might employ varying strategies on a trial-by-trial basis, even within a single-day skill acquisition period. Second, the direction of errors for both obstacles predominantly involved reducing foot clearance for most participants, which may limit the richness of our dataset in accurately estimating the presence of interference using the current model. Due to participants’ physical challenges with increasing foot clearance, we could not incorporate opposing directions of error between two heights of obstacles for most participants. This limitation might have affected our ability to detect interference accurately. Despite these challenges, previous studies have primarily examined the effect of interference on motor retention in people with PD (31,35,47,51,52), but the dynamics of learning multiple skills during skill acquisition have not been systematically studied in this population. Our results offer novel insights into how individuals may experience varying magnitudes of interference and that even within an individual, the influence of interference may fluctuate on a trial-by-trial basis. These dynamics can enhance our understanding of how interference affects other aspects of motor learning, such as motor retention. Future studies should continue to explore these dynamics to better inform the task structures for physical rehabilitation.

Further, while online performance improvement is intact in people with PD, as hypothesized, we also found no significant differences in overnight retention in people with PD compared to age-matched controls. The lack of group differences in overnight retention was inconsistent with the impaired retention found in the previous study on locomotor learning using exergaming (34), yet our results were consistent with other studies investigating balance learning (41,43,49). The exergaming study included participants who scored 24 or higher for the mini-mental state examination (MMSE) (34). This scale often finds 26 as a target with mild cognitive impairments or mild dementia for people with PD (81–83). Including individuals who scored MMSE lower than 26 may indicate that their participants were more cognitively impaired than ours. This suggests that people with PD, particularly those with normal cognition, have intact locomotor retention abilities comparable to the age-matched adults. Despite the disruption of the cortico-striatal circuits from the time of diagnosis in PD, compensatory effects from other brain regions such as prefrontal cortex and cerebellum also come into play to maintain motor functions in people with PD (70,84,85). For instance, people with PD at H&Y stage 2 exhibit increased activity in the prefrontal cortex during obstacle negotiation, suggesting that the prefrontal cortex may compensate for attention and motor deficits (84). During retention, other brain regions, such as the prefrontal cortex (74) or cerebellum (70,85), may further compensate to allow people with PD to perform the skill with greater effort yet still exhibit intact retention. Further research is warranted to explore the neural mechanisms, specifically the contribution of compensatory effects, underlying intact motor retention in people with early stages of PD without cognitive impairments to inform the design of physical rehabilitation for people with PD.

The distinct associations between online performance improvement and overnight retention of locomotor skills with learning parameters may suggest that how individuals learn locomotor skills influences retention over varying timeframes. Moreover, these associations were stronger in the control group than in the PD group, indicating a diminished impact of how individuals acquire locomotor skills on the retention environment in people with PD. In contrast to previous studies linking learning rates to long-term retention (86–88), our findings in controls show associations where slower learning rates are associated with better online performance improvement while low interference is associated with better overnight retention. This discrepancy with previous studies may arise from the complexity of locomotor tasks. We investigated locomotor skill learning, which requires precise foot control and coordination with other body parts, such as the trunk, while previous studies focused on finger or arm movements. For locomotor skill learning, slower learning might facilitate a better understanding of tasks, enhancing online performance improvement without feedback.

Conversely, faster learners might adopt incomplete strategies early on, worsening performance when performance feedback is removed. Regarding contextual interference, less interference while learning multiple skills in a randomized order might allow individuals to develop more suitable strategies for acquiring different skills simultaneously. While these strategies may not lead to immediate benefits during skill acquisition, they may contribute to better retention of the locomotor skills overnight. This highlights the importance of optimal challenge points (47) for enhancing long-term retention. Additionally, consistent with our previous study in young adults (69), both groups exhibited a significant relationship between performance at the end of skill acquisition and overnight retention. This consistency suggests that the final performance level achieved during initial locomotor skill acquisition is a robust indicator of long-term retention across different populations. These differential patterns underscore the importance of tailored rehabilitation strategies. Future research can explore motor learning involving whole-body movements and trial-by-trial interference to understand individual differences in locomotor skill learning.

There are notable limitations of the study. First, our sample size was relatively small, particularly in the PD group. A larger sample size in the future would enhance the generalizability of our findings and provide better insights into how individual differences in learning performance influence retention. Second, while the population parameter for learning rate was biased toward faster learning in people with PD, the group difference credible intervals spanned over zero, indicating an absence of definitive evidence. This likely reflects the small sample size in our current study and highlights the need for future research with larger samples to confirm these findings. Despite these limitations, our study provides valuable insights into how early stages of PD without cognitive impairments influence different aspects of locomotor skill learning compared to controls.

## Conclusion

In this study, we demonstrated that people with early stages of PD acquire locomotor skills more rapidly than age-matched controls, but there is limited evidence of differences in interference between the two groups while learning multiple objectives. However, both groups exhibited comparable performance improvement during skill acquisition and retention after 24 hours. Notably, our study identified associations between retention and learning parameters in the control group: learning rate was associated with online performance improvement, while the degree of interference was linked to overnight retention. Moreover, regardless of the group, final performance at the end of skill acquisition was a significant indication for overnight retention. These findings highlight that people with early stages of PD retain the capacity for complex locomotor skill learning, but individual differences in the learning process may differentially influence online performance improvement and overnight retention.

## References

1. Braak H, Del Tredici K, Rüb U, de Vos RAI, Jansen Steur ENH, Braak E. Staging of brain pathology related to sporadic Parkinson’s disease. Neurobiol Aging. 2003;24(2):197–211.

2. Hawkes CH, Del Tredici K, Braak H. A timeline for Parkinson’s disease. Parkinsonism Relat Disord. 2010 Feb;16(2):79–84.

3. Peterson DS, Horak FB. Neural Control of Walking in People with Parkinsonism. Physiol Bethesda Md. 2016 Mar;31(2):95–107.

4. Armstrong MJ, Okun MS. Diagnosis and Treatment of Parkinson Disease: A Review. JAMA. 2020 Feb 11;323(6):548–60.

5. Dirnberger G, Jahanshahi M. Executive dysfunction in Parkinson’s disease: A review. J Neuropsychol. 2013 Sep 1;7(2):193–224.

6. Goldman JG, Vernaleo BA, Camicioli R, Dahodwala N, Dobkin RD, Ellis T, et al. Cognitive impairment in Parkinson’s disease: a report from a multidisciplinary symposium on unmet needs and future directions to maintain cognitive health. Npj Park Dis. 2018 Jun 26;4(1):1–11.

7. Dagan M, Herman T, Bernad-Elazari H, Gazit E, Maidan I, Giladi N, et al. Dopaminergic therapy and prefrontal activation during walking in individuals with Parkinson’s disease: does the levodopa overdose hypothesis extend to gait? J Neurol. 2021 Feb;268(2):658–68.

8. Fisher BE, Li Q, Nacca A, Salem GJ, Song J, Yip J, et al. Treadmill exercise elevates striatal dopamine D2 receptor binding potential in patients with early Parkinson’s disease: NeuroReport. 2013 Jul;24(10):509–14.

9. Petzinger GM, Fisher BE, Van Leeuwen JE, Vukovic M, Akopian G, Meshul CK, et al. Enhancing neuroplasticity in the basal ganglia: the role of exercise in Parkinson’s disease. Mov Disord Off J Mov Disord Soc. 2010;25 Suppl 1:S141–145.

10. Petzinger GM, Fisher BE, McEwen S, Beeler JA, Walsh JP, Jakowec MW. Exercise-enhanced neuroplasticity targeting motor and cognitive circuitry in Parkinson’s disease. Lancet Neurol. 2013 Jul;12(7):716–26.

11. Gisbert R, Schenkman M. Physical Therapist Interventions for Parkinson Disease. Phys Ther. 2015 Mar 1;95(3):299–305.

12. Levin MF, Demers M. Motor learning in neurological rehabilitation. Disabil Rehabil. 2020 Apr 22;1–9.

13. Wolpert DM, Diedrichsen J, Flanagan JR. Principles of sensorimotor learning. Nat Rev Neurosci. 2011 Dec;12(12):739–51.

14. Spampinato D, Celnik P. Multiple Motor Learning Processes in Humans: Defining Their Neurophysiological Bases. The Neuroscientist. 2020 Jul 25;27(3):246.

15. Krakauer JW, Hadjiosif AM, Xu J, Wong AL, Haith AM. Motor Learning. Compr Physiol. 2019 Mar 14;9(2):613–63.

16. Huang VS, Haith A, Mazzoni P, Krakauer JW. Rethinking Motor Learning and Savings in Adaptation Paradigms: Model-Free Memory for Successful Actions Combines with Internal Models. Neuron. 2011 May 26;70(4):787–801.

17. Mazzoni P, Krakauer JW. An implicit plan overrides an explicit strategy during visuomotor adaptation. J Neurosci Off J Soc Neurosci. 2006 Apr 5;26(14):3642–5.

18. Tseng Y weng, Diedrichsen J, Krakauer JW, Shadmehr R, Bastian AJ. Sensory Prediction Errors Drive Cerebellum-Dependent Adaptation of Reaching. J Neurophysiol. 2007 Jul 1;98(1):54–62.

19. Therrien AS, Wolpert DM, Bastian AJ. Effective reinforcement learning following cerebellar damage requires a balance between exploration and motor noise. Brain. 2016 Jan;139(1):101–14.

20. Galea JM, Mallia E, Rothwell J, Diedrichsen J. The dissociable effects of punishment and reward on motor learning. Nat Neurosci. 2015 Apr;18(4):597–602.

21. Izawa J, Shadmehr R. Learning from Sensory and Reward Prediction Errors during Motor Adaptation. Körding K, editor. PLoS Comput Biol. 2011 Mar 10;7(3):e1002012.

22. Reinkensmeyer DJ, Burdet E, Casadio M, Krakauer JW, Kwakkel G, Lang CE, et al. Computational neurorehabilitation: modeling plasticity and learning to predict recovery. J Neuroengineering Rehabil. 2016 Apr 30;13(1):42.

23. Jax SA, Rosenbaum DA. Hand path priming in manual obstacle avoidance: evidence that the dorsal stream does not only control visually guided actions in real time. J Exp Psychol Hum Percept Perform. 2007 Apr;33(2):425–41.

24. Diedrichsen J, White O, Newman D, Lally N. Use-Dependent and Error-Based Learning of Motor Behaviors. J Neurosci. 2010 Apr 14;30(15):5159–66.

25. Popp NJ, Yokoi A, Gribble PL, Diedrichsen J. The effect of instruction on motor skill learning. J Neurophysiol. 2020 Nov 1;124(5):1449–57.

26. Meier C, Frank C, Gröben B, Schack T. Verbal Instructions and Motor Learning: How Analogy and Explicit Instructions Influence the Development of Mental Representations and Tennis Serve Performance. Front Psychol. 2020 Feb 7;11:2.

27. Wulf G, Höß M, Prinz W. Instructions for Motor Learning: Differential Effects of Internal Versus External Focus of Attention. J Mot Behav. 1998 Jun 1;30(2):169–79.

28. Yamada C, Itaguchi Y, Fukuzawa K. Effects of the amount of practice and time interval between practice sessions on the retention of internal models. PLoS ONE. 2019 Apr 16;14(4):e0215331.

29. Shadmehr R. Generalization as a behavioral window to the neural mechanisms of learning internal models. Hum Mov Sci. 2004 Nov;23(5):543–68.

30. Krakauer JW, Mazzoni P, Ghazizadeh A, Ravindran R, Shadmehr R. Generalization of Motor Learning Depends on the History of Prior Action. PLOS Biol. 2006 Sep 12;4(10):e316.

31. Shea JB, Morgan RL. Contextual interference effects on the acquisition, retention, and transfer of a motor skill. J Exp Psychol [Hum Learn]. 1979;5(2):179–87.

32. van Hedel HJA, Waldvogel D, Dietz V. Learning a high-precision locomotor task in patients with Parkinson’s disease. Mov Disord Off J Mov Disord Soc. 2006 Mar;21(3):406–11.

33. Michel J, Benninger D, Dietz V, van Hedel HJA. Obstacle stepping in patients with Parkinson’s disease. Complexity does influence performance. J Neurol. 2009 Mar;256(3):457–63.

34. Mendes FA dos S, Pompeu JE, Lobo AM, da Silva KG, Oliveira T de P, Zomignani AP, et al. Motor learning, retention and transfer after virtual-reality-based training in Parkinson’s disease – effect of motor and cognitive demands of games: a longitudinal, controlled clinical study. Physiotherapy. 2012 Sep;98(3):217–23.

35. Lin CH (Janice), Sullivan KJ, Wu AD, Kantak S, Winstein CJ. Effect of Task Practice Order on Motor Skill Learning in Adults With Parkinson Disease: A Pilot Study. Phys Ther. 2007 Sep 1;87(9):1120–31.

36. Soliveri P, Brown RG, Jahanshahi M, Caraceni T, Marsden CD. Learning manual pursuit tracking skills in patients with Parkinson’s disease. Brain J Neurol. 1997 Aug;120 (Pt 8):1325–37.

37. Swinnen SP, Steyvers M, Van Den Bergh L, Stelmach GE. Motor learning and Parkinson’s disease: refinement of within-limb and between-limb coordination as a result of practice. Behav Brain Res. 2000 Jun 15;111(1):45–59.

38. Leow LA, Loftus AM, Hammond GR. Impaired savings despite intact initial learning of motor adaptation in Parkinson’s disease. Exp Brain Res. 2012 Apr 1;218(2):295–304.

39. Bédard P, Sanes JN. Basal ganglia-dependent processes in recalling learned visual-motor adaptations. Exp Brain Res. 2011 Mar;209(3):385–93.

40. Leow LA, de Rugy A, Loftus AM, Hammond G. Different mechanisms contributing to savings and anterograde interference are impaired in Parkinson’s disease. Front Hum Neurosci. 2013;7:55.

41. Van Ooteghem K, Frank JS, Horak FB. Postural motor learning in Parkinson’s disease: The effect of practice on continuous compensatory postural regulation. Gait Posture. 2017;57:299–304.

42. Nemanich ST, Earhart GM. Reduced after-effects following podokinetic adaptation in people with Parkinson’s disease and freezing of gait. Parkinsonism Relat Disord. 2016 Jan;22:93–7.

43. Jessop RT, Horowicz C, Dibble LE. Motor learning and Parkinson disease: Refinement of movement velocity and endpoint excursion in a limits of stability balance task. Neurorehabil Neural Repair. 2006 Dec;20(4):459–67.

44. Pendt LK, Maurer H, Müller H. The Influence of Movement Initiation Deficits on the Quantification of Retention in Parkinson’s Disease. Front Hum Neurosci. 2012 Aug 1;6:226.

45. Pendt LK, Reuter I, Müller H. Motor Skill Learning, Retention, and Control Deficits in Parkinson’s Disease. PLOS ONE. 2011 Jul 8;6(7):e21669.

46. Behrman AL, Cauraugh JH, Light KE. Practice as an intervention to improve speeded motor performance and motor learning in Parkinson’s disease. J Neurol Sci. 2000 Mar 15;174(2):127–36.

47. Onla-or S, Winstein CJ. Determining the optimal challenge point for motor skill learning in adults with moderately severe Parkinson’s disease. Neurorehabil Neural Repair. 2008 Aug;22(4):385–95.

48. Lee YY, Winstein CJ, Gordon J, Petzinger GM, Zelinski EM, Fisher BE. Context-Dependent Learning in People With Parkinson’s Disease. J Mot Behav. 2015 Sep 16;1–9.

49. Smiley-Oyen AL, Lowry KA, Emerson QR. Learning and retention of movement sequences in Parkinson’s disease. Mov Disord Off J Mov Disord Soc. 2006 Aug;21(8):1078–87.

50. Lee YY, Tai CH, Fisher BE. Context-Dependent Behavior in Parkinson’s Disease With Freezing of Gait. Neurorehabil Neural Repair. 2019 Dec 1;33(12):1040–9.

51. Sidaway B, Ala B, Baughman K, Glidden J, Cowie S, Peabody A, et al. Contextual Interference Can Facilitate Motor Learning in Older Adults and in Individuals With Parkinson’s Disease. J Mot Behav. 2016 Nov 1;48(6):509–18.

52. Lahlou S, Gabitov E, Owen L, Shohamy D, Sharp M. Preserved motor memory in Parkinson’s disease. Neuropsychologia. 2022 Mar 12;167:108161.

53. Roemmich RT, Nocera JR, Stegemöller EL, Hassan A, Okun MS, Hass CJ. Locomotor adaptation and locomotor adaptive learning in Parkinson’s disease and normal aging. Clin Neurophysiol. 2014 Feb 1;125(2):313–9.

54. Corzani M, Ferrari A, Ginis P, Nieuwboer A, Chiari L. Motor Adaptation in Parkinson’s Disease During Prolonged Walking in Response to Corrective Acoustic Messages. Front Aging Neurosci. 2019;11:265.

55. Wu T, Hallett M. The cerebellum in Parkinson’s disease. Brain. 2013 Mar;136(3):696–709.

56. Nagano-Saito A, Leyton M, Monchi O, Goldberg YK, He Y, Dagher A. Dopamine depletion impairs frontostriatal functional connectivity during a set-shifting task. J Neurosci Off J Soc Neurosci. 2008 Apr 2;28(14):3697–706.

57. Cho J, Duke D, Manzino L, Sonsalla PK, West MO. Dopamine depletion causes fragmented clustering of neurons in the sensorimotor striatum: evidence of lasting reorganization of corticostriatal input. J Comp Neurol. 2002 Oct 7;452(1):24–37.

58. Turner RS, Desmurget M. Basal Ganglia Contributions to Motor Control: A Vigorous Tutor. Curr Opin Neurobiol. 2010 Dec;20(6):704–16.

59. Doyon J, Bellec P, Amsel R, Penhune V, Monchi O, Carrier J, et al. Contributions of the basal ganglia and functionally related brain structures to motor learning. Behav Brain Res. 2009 Apr 12;199(1):61–75.

60. Litvan I, Goldman JG, Tröster AI, Schmand BA, Weintraub D, Petersen RC, et al. Diagnostic Criteria for Mild Cognitive Impairment in Parkinson’s Disease: Movement Disorder Society Task Force Guidelines. Mov Disord Off J Mov Disord Soc. 2012 Mar;27(3):349–56.

61. Smith MA, Ghazizadeh A, Shadmehr R. Interacting Adaptive Processes with Different Timescales Underlie Short-Term Motor Learning. PLOS Biol. 2006 May 23;4(6):e179.

62. Browne WJ, Draper D. A comparison of Bayesian and likelihood-based methods for fitting multilevel models. Bayesian Anal. 2006 Sep;1(3):473–514.

63. Kruschke J. Doing Bayesian Data Analysis: A Tutorial with R, JAGS, and Stan. 2nd edition. Boston: Academic Press; 2014. 776 p.

64. Gelman A, Rubin DB. Inference from Iterative Simulation Using Multiple Sequences. Stat Sci. 1992;7(4):457–72.

65. Vehtari A, Gelman A, Simpson D, Carpenter B, Bürkner PC. Rank-Normalization, Folding, and Localization: An Improved R for Assessing Convergence of MCMC. Bayesian Anal. 2021 Jan;1(1):1–28.

66. Makowski D, Ben-Shachar MS, Chen SHA, Lüdecke D. Indices of Effect Existence and Significance in the Bayesian Framework. Front Psychol. 2019 Dec 10;10:2767.

67. Makowski D, Ben-Shachar MS, Lüdecke D. bayestestR: Describing Effects and their Uncertainty, Existence and Significance within the Bayesian Framework. J Open Source Softw. 2019 Aug 13;4(40):1541.

68. Kruschke JK. Bayesian estimation supersedes the t test. J Exp Psychol Gen. 2013 May;142(2):573– 603.

69. Kim A, Schweighofer N, Finley JM. Locomotor skill acquisition in virtual reality shows sustained transfer to the real world. J NeuroEngineering Rehabil. 2019 Sep 14;16(1):113.

70. Mentis MJ, Dhawan V, Nakamura T, Ghilardi MF, Feigin A, Edwards C, et al. Enhancement of brain activation during trial-and-error sequence learning in early PD. Neurology. 2003 Feb 25;60(4):612–9.

71. Yu H, Sternad D, Corcos DM, Vaillancourt DE. Role of Hyperactive Cerebellum and Motor Cortex in Parkinson’s Disease. NeuroImage. 2007 Mar;35(1):222–33.

72. Xu S, He XW, Zhao R, Chen W, Qin Z, Zhang J, et al. Cerebellar functional abnormalities in early stage drug-naïve and medicated Parkinson’s disease. J Neurol. 2019 Jul;266(7):1578–87.

73. Solstrand Dahlberg L, Lungu O, Doyon J. Cerebellar Contribution to Motor and Non-motor Functions in Parkinson’s Disease: A Meta-Analysis of fMRI Findings. Front Neurol. 2020;11:127.

74. Martin JA, Zimmermann N, Scheef L, Jankowski J, Paus S, Schild HH, et al. Disentangling motor planning and motor execution in unmedicated de novo Parkinson’s disease patients: An fMRI study. NeuroImage Clin. 2019 Mar 19;22:101784.

75. Wolpert DM, Miall RC, Kawato M. Internal models in the cerebellum. Trends Cogn Sci. 1998 Sep;2(9):338–47.

76. Popa LS, Ebner TJ. Cerebellum, Predictions and Errors. Front Cell Neurosci. 2018;12:524.

77. Morton SM, Bastian AJ. Cerebellar contributions to locomotor adaptations during splitbelt treadmill walking. J Neurosci Off J Soc Neurosci. 2006 Sep 6;26(36):9107–16.

78. Werheid K, Zysset S, Müller A, Reuter M, von Cramon DY. Rule learning in a serial reaction time task: an fMRI study on patients with early Parkinson’s disease. Brain Res Cogn Brain Res. 2003 Apr;16(2):273–84.

79. Bédard P, Sanes JN. On a basal ganglia role in learning and rehearsing visual–motor associations. NeuroImage. 2009 Oct 1;47(4):1701–10.

80. Morton SM, Bastian AJ. Cerebellar Control of Balance and Locomotion. The Neuroscientist. 2004 Jun 1;10(3):247–59.

81. Hoops S, Nazem S, Siderowf AD, Duda JE, Xie SX, Stern MB, et al. Validity of the MoCA and MMSE in the detection of MCI and dementia in Parkinson disease. Neurology. 2009 Nov 24;73(21):1738–45.

82. Kaszás B, Kovács N, Balás I, Kállai J, Aschermann Z, Kerekes Z, et al. Sensitivity and specificity of Addenbrooke’s Cognitive Examination, Mattis Dementia Rating Scale, Frontal Assessment Battery and Mini Mental State Examination for diagnosing dementia in Parkinson’s disease. Parkinsonism Relat Disord. 2012 Jun;18(5):553–6.

83. Dujardin K, Dubois B, Tison F, Durif F, Bourdeix I, Péré JJ, et al. Parkinson’s disease dementia can be easily detected in routine clinical practice. Mov Disord Off J Mov Disord Soc. 2010 Dec 15;25(16):2769–76.

84. Assad M, Galperin I, Giladi N, Mirelman A, Hausdorff JM, Maidan I. Disease severity and prefrontal cortex activation during obstacle negotiation among patients with Parkinson’s disease: Is it all as expected? Parkinsonism Relat Disord. 2022 Aug 1;101:20–6.

85. Wu T, Hallett M. A functional MRI study of automatic movements in patients with Parkinson’s disease. Brain J Neurol. 2005 Oct;128(Pt 10):2250–9.

86. Wadden KP, Asis KD, Mang CS, Neva JL, Peters S, Lakhani B, et al. Predicting Motor Sequence Learning in Individuals With Chronic Stroke. Neurorehabil Neural Repair. 2017;31(1):95–104.

87. Schaefer SY, Duff K. Rapid Responsiveness to Practice Predicts Longer-Term Retention of Upper Extremity Motor Skill in Non-Demented Older Adults. Front Aging Neurosci. 2015 Nov 18;7:214.

88. Park H, Schweighofer N. Nonlinear mixed-effects model reveals a distinction between learning and performance in intensive reach training post-stroke. J Neuroengineering Rehabil. 2017 Mar 17;14(1):21.

